# Mixed Effects Association of Single Cells Identifies an Expanded Th1-Skewed Cytotoxic Effector CD4+ T Cell Subset in Rheumatoid Arthritis

**DOI:** 10.1101/172403

**Authors:** Chamith Y. Fonseka, Deepak A. Rao, Nikola C. Teslovich, Susan K. Hannes, Kamil Slowikowsi, Michael F. Gurish, Laura T. Donlin, Michael E. Weinblatt, Elena M. Massarotti, Jonathan S. Coblyn, Simon M. Helfgott, Derrick J. Todd, Vivian P. Bykerk, Elizabeth W. Karlson, Joerg Ermann, Yvonne C. Lee, Michael B. Brenner, Soumya Raychaudhuri

## Abstract

High dimensional single-cell analyses have dramatically improved the ability to resolve complex mixtures of cells from human disease samples; however, identifying disease-associated cell types or cell states in patient samples remains challenging due to technical and inter-individual variation. Here we present Mixed effects modeling of Associations of Single Cells (MASC), a novel reverse single cell association strategy for testing whether case-control status influences the membership of single cells in any of multiple cellular subsets while accounting for technical confounds and biological variation. Applying MASC to mass cytometry analyses of CD4+ T cells from blood of rheumatoid arthritis (RA) patients and controls revealed a significantly expanded population of CD4+ T cells, identified as CD27- HLA-DR+ effector memory cells, in RA patients (OR = 1.7; p = 1.1 × 10^−3^). The frequency of CD27- HLA-DR+ cells was similarly elevated in blood samples from a second RA patient cohort, and CD27- HLA-DR+ cell frequency decreased in RA patients who respond to immunosuppressive therapy. Compared to peripheral blood, synovial fluid and synovial tissue samples from RA patients contained ∼5-fold higher frequencies of CD27- HLA-DR+ cells, which comprised ∼10% of synovial CD4+ T cells. We find that CD27- HLA-DR+ cells are abundant producers of IFN-γ and also express perforin and granzyme A at elevated levels. Thus MASC identified the expansion of a unique Th1 skewed effector T cell population with cytotoxic capacity in RA. We propose that MASC is a broadly applicable method to identify disease-associated cell populations in high-dimensional single cell data.

**One Sentence Summary:** Mixed-effects regression of single cells identifies a cytotoxic Th1-like CD4+ T cell subset while accounting for inter-individual and technical variation.

## Main Text

## Introduction

The advance of single cell technologies has enabled investigators to resolve cellular heterogeneity with unprecedented resolution. Single cell assays have been particularly useful in the study of the immune system, in which diverse cell populations often consisting of rare and transitional cell states may play an important role *(1)*. Application of single cell transcriptomic and cytometric assays in a case-control study has the potential to reveal expanded pathogenic cell populations in immune-mediated diseases.

Rheumatoid arthritis (RA) is a chronic, systemic disease affecting 0.5-1% of the adult population, making it one of the most common autoimmune disorders worldwide *(2)*. RA is triggered by environmental and genetic risk factors, leading to activation of autoreactive T cells and B cells that mediate an autoimmune response directed at the joints *(3, 4)*. CD4+ T cells have been strongly implicated in RA pathogenesis *(5, 6)*. For one, the strongest genetic association to RA is with the *HLA-DRB1* gene within the MHC; these polymorphisms affect the range of antigens that MHCII molecules can bind and present in order to activate CD4+ T cells *(3, 7, 8)*. Furthermore, many RA risk alleles outside of the MHC locus also lie in pathways important for CD4+ T cell activation, differentiation into effector (T_eff_) and regulatory (T_reg_) subsets, and maintenance of subset identity *(4, 8-12)*. Defining the precise CD4+ T cell subsets that are expanded or dysregulated in RA patients is critical to deciphering pathogenesis. Such cell populations may be enriched in antigen-specific T cells and may aid in discovery of dominant disease-associated autoantigens. In addition, these populations may directly carry out pathologic effectors functions that can be targeted therapeutically *(13)*.

For many autoimmune diseases, directly assaying affected tissues is difficult because samples are only available through invasive procedures. Instead, querying peripheral blood for altered immune cell populations is a rapidly scalable strategy that achieves larger sample sizes and allows for serial monitoring. Flow cytometric studies have identified alterations in specific circulating T cell subsets in RA patients, including an increased frequency of CD28- CD4+ T cells *(14-16)*; however, the expansion of CD28- T cells represents one of the relatively few T cell alterations that has been reproducibly detected by multiple groups. Limited reproducibility may be the consequence of differences in clinical cohorts, small sample sizes, methodologic variability, and use of limited, idiosyncratic combinations of phenotypic markers*(17)*.

The advent of mass cytometry now allows for relatively broad assessment of circulating immune cell populations with >30 markers *(18)*, enabling detailed, multiparametric characterization of lymphocyte subsets. This technology provides the potential to define and quantify lymphocyte subsets at high resolution using multiple markers. While rapid progress has been made in defining individual cellular populations in an unbiased fashion *(19-23)*, a key challenge that remains is establishing methods to identify cell populations associated with a disease. In particular, inter-individual variation and technical variation influence cell population frequencies and need to be accounted for in an association framework.

Here we describe a robust statistical method to test for disease associations with single cell data called MASC (Mixed effect modeling of Associations of Single Cells), which tests at the single cell level the association between population clusters and disease status. It is a ‘reverse’ association strategy where the case-control status is an independent variable, rather than the dependent variable. We have applied MASC to identify T cell subsets associated with RA in a mass cytometry case-control immunophenotyping dataset that we generated focused on CD4+ T cells. This high-dimensional analysis enabled us to identify disease-specific changes in canonical as well as non-canonical CD4+ T cell populations using a panel of 32 markers to reveal cell lineage, activation, and function *(24, 25)*. Using MASC, we identify a population of memory CD4+ T cells, characterized as CD27- HLA-DR+, that is expanded in the circulation of RA patients. Further, we find that CD27- HLA-DR+ T cells are enriched within inflamed RA joints, rapidly produce IFN-γ and cytolytic factors, and contract with successful treatment of RA.

## Results

### Statistical and computational strategy

Single cell association studies, with either mass cytometry or single-cell RNA-seq, require statistical strategies robust to inter-individual donor variability and technical effects that can skew cell subset estimates. For example, mass cytometry studies need to control for variability in machine sensitivity, reagent staining, and sample handling that can lead to batch effects. Inter-individual differences can lead to real shifts in cell population frequencies at baseline, while technical effects can lead to apparent shifts in cell population frequencies. MASC test for case-control differences while accounting for technical and inter-individual variation. We acquired single cell data from individual case-control samples (**Figure 1**), and then applied MASC after stringent quality controls (see **Methods**).

**Figure 1:**
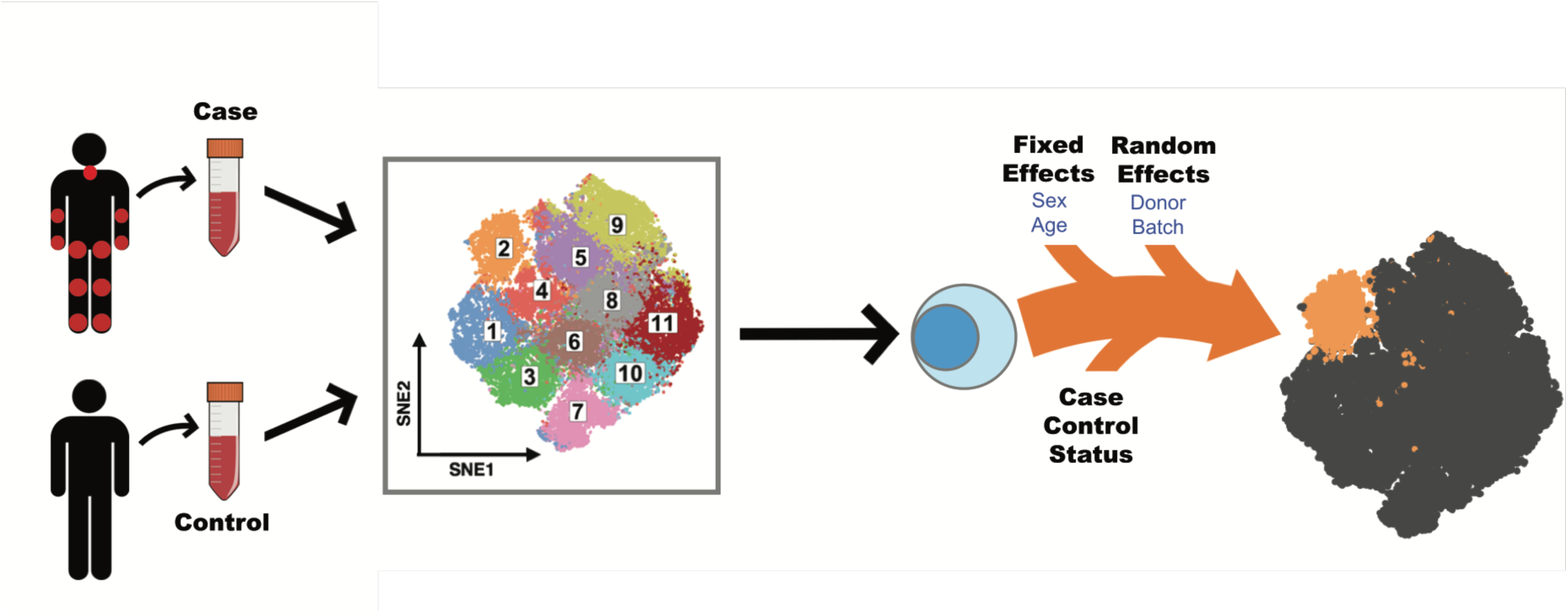
Mixed effect modeling of Associations of Single Cells (MASC) Overview. Single cell transcriptomics or proteomics are used to assay samples from cases and controls, such as immunoprofiling of peripheral blood. The data is then clustered to define populations of similar cells. Mixed-effect logistic regression is used to predict individual cell membership in previously defined populations. The addition of a case-control term to the regression model allows the user to identify populations for which case-control status is significantly associated.

First, we objectively define unbiased subsets of cells using an equal numbers of random cells from each sample so that they contributed equally to the subsequent analyses. Here we used DensVM *(26)* to cluster mass cytometry data, but recognize the availability of alternative methodologies. Next, we used MASC, a reverse association strategy, using single cell logistic mixed-effect modeling to test individual subsets for association. Here, we predict the subset membership of each cell based upon fixed effects (e.g. sex) and random effects (e.g. batch, donor) in a null model. We then measure the improvement in model fit when a fixed effect term for the case-control status of the sample is included with a likelihood ratio test. This framework allowed us to evaluate the significance and effect size of the case-control association for each subset while controlling for inter-individual and technical variability.

To ensure that MASC was robust to statistical inflation, we ran it 10,000 times after permuting case-control labels using the dataset described below to obviate almost any case-control association that might be present. We then recorded the reported p-values for each cluster. This approach demonstrated that MASC has only a modestly inflated type I error rate for the 19 clusters in the dataset with 6.5% of trials obtaining p<0.05 (**Figure S2A**). Importantly, we found that including both donor and batch random effects was critical; eliminating random effects in the model led to highly inflated p-values with 66.1% of trials obtaining p<0.05 (**Figure S2B**).

As an alternative and frequently used strategy, we also tested a simple binomial test for case-control association. This approach is limited in our view as it fails to model donor specific and technical effects. Unsurprisingly, we found that this commonly used approach produced highly inflated results in comparison to MASC with 65.7% of trials obtaining p<0.05 (**Figure S2C**).

### Experimental strategy

Here, we applied this method to mass cytometric analysis of CD4+ T cells obtained from peripheral blood of patients with established RA (**Table 1**). We purified memory CD4+ T cells from RA patients (cases) and non-inflammatory controls and either (1) rested the cells for 24 hours or (2) stimulated the cells with anti-CD3/anti-CD28 beads for 24 hours. We then analyzed the cells using a 32-marker mass cytometry panel that included 22 markers of lineage, activation, and function (**Supplementary Table 1**). This design allowed us to interrogate immune states across cases and controls and also capture stimulation-dependent changes. After stringent quality control measures, we analyzed a total of 26 cases and 26 controls. We randomly sampled 1000 cells from each of the 52 samples so that each sample contributed equally to the analysis. We then used Barnes-Hut SNE algorithm to project the data altogether so that all cells from all samples were projected into the same two dimensions, and then applied DensVM to assign cells to subsets by their similarity of marker expression *(26)*.

**Table 1:**
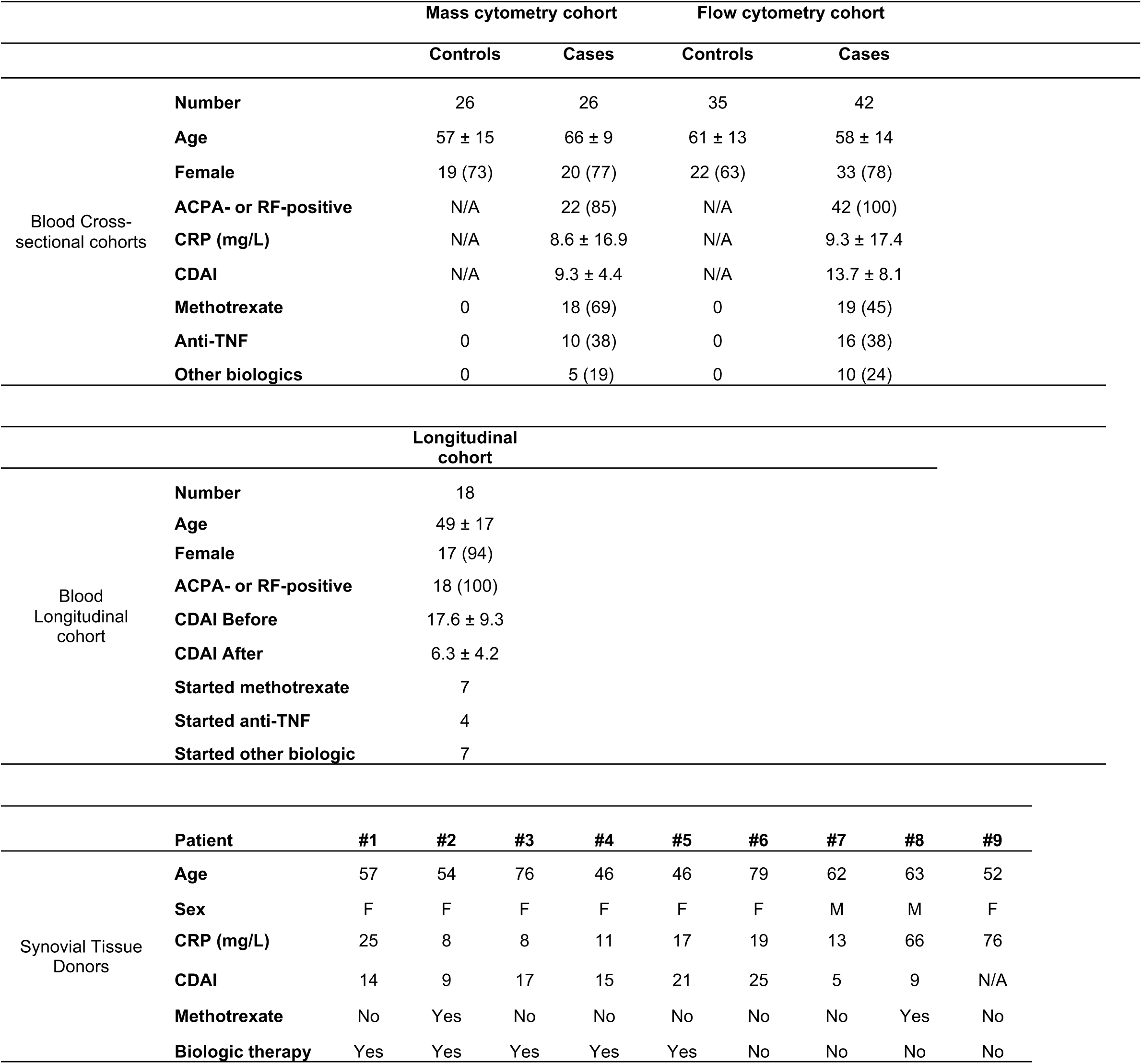
Clinical characteristics of patient samples used. Average ± SD is shown. Parentheses indicate percentages. CRP: C-reactive protein. CDAI: clinical disease activity index. Other biologics include rituximab, tofacitinib, and abatacept.

### Mass Cytometry Reveals The Landscape of CD4+ Memory T Cell Subsets

We observed substantial diversity among resting CD4+ memory T cells, consistent with previous reports demonstrating the breadth of phenotypes in CD8+ T cells and CD4+ T cells *(27-29)*. We identified 19 distinct subsets in resting (*R*) memory CD4+ T cells (**Figure 2A**). Central memory T cells (T_CM_) segregated from effector memory T cells (T_EM_) by the expression of CD62L (**Figure 2B**). Five subsets (subsets 1 – 5) of central memory T (T_CM_) cells, all expressing CD62L, varied in the expression of CD27 and CD38, highlighting the heterogeneity within the T_CM_ compartment (**Figure 2E**). We were able to identify two T_h_1 subsets (subsets 8 and 12) as well as two T_reg_ subsets (subsets 7 and 11). While both T_reg_ subsets expressed high levels of CD25 and FoxP3, subset 11 also expressed HLA-DR, reflecting a known diversity among T_reg_ populations in humans *(30)*.

**Figure 2:**
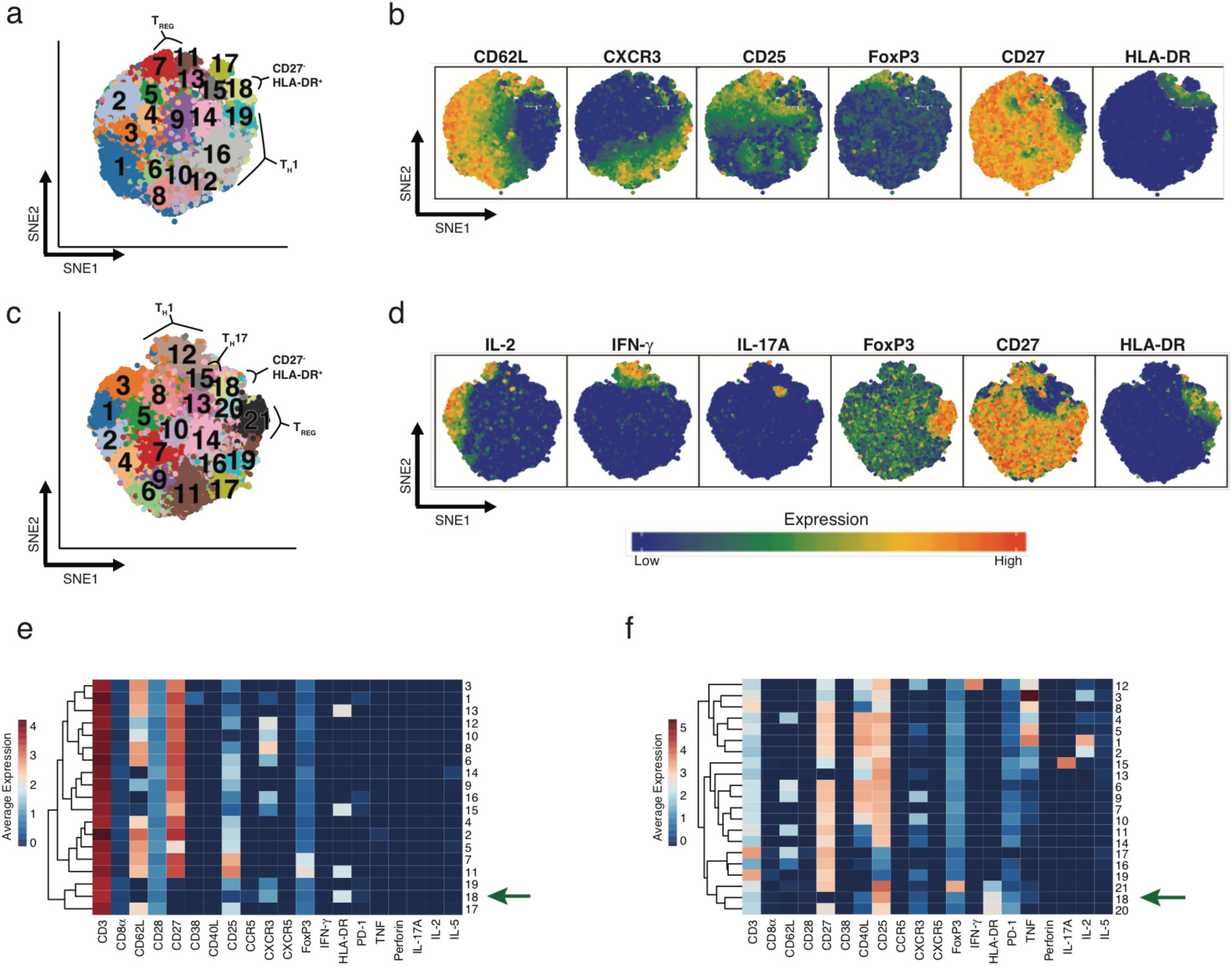
Diversity of CD4^+^ memory T cells before and after stimulation. (a) SNE projection of 50,000 resting CD4+ memory T cells sampled equally from RA patients (n=24) and controls (n=26). DensVM identified 19 populations in this dataset. (b) Same SNE projections as in (a) colored by the expression of the marker labeled above each plot. (c) SNE projection of 52,000 CD4^+^ stimulated memory T cells sampled equally from RA patients (n=26) and controls (n=26). Cells were stimulated for 24 hours with anti-CD3/anti-CD28 beads. (d) Same SNE projections as in (c) colored by the expression of the marker labeled above each plot. (e) Heatmap showing mean expression of indicated markers across the 19 populations found in resting cells. (f) Heatmap showing mean expression of indicated markers across the 21 populations found after stimulation. Protein expression data are shown after arcsinh transformation.

When applied to the stimulated CD4+ T cell data, DensVM identified 21 subsets (**Figure 2C**). As expected, certain activation markers, like CD25 and CD40L, were broadly expressed across most subsets after stimulation (**Figure 2D**). Mass cytometry robustly detected cytokine production from stimulated CD4+ memory T cells (**Figure 2D, F**). Activated effector cells identified by the production of IL-2 were separated into three groups (subsets 1 – 3) by relative expression of TNF. Cytokine expression after stimulation also improved the ability to resolve certain CD4+ T effector subsets, such as T_h_17 cells (subset 15) and T_h_1 cells (subset 12). However, we were unable to resolve the T_reg_ subtypes that were observed in the resting T cells; after stimulation, all CD25+ FoxP3+ cells were grouped together (cluster 21). The set of cells that did not activate after stimulation (subsets 16, 17, and 19) were easily identified by the lack of expression of activation markers such as CD25, CD40L and cytokines and the retention of high CD3 expression.

### Identifying Populations that are Enriched or Depleted in RA Samples

We next sought to identify subsets that are significantly overrepresented or underrepresented by cells from cases. The percentage of RA cells in each subset ranged from 36.7% to 63.6% in resting state, and 38.9% to 65.7% in the stimulated state (**Figure S1A,B**). When we formally tested whether they were statistically significant accounting for subject-specific and batch-specific random effects with MASC (**Methods**), we observed three populations with significantly altered proportions of cells from cases in the resting T cell data. Most notably, we observed enrichment in subset 18 (p = 5.9 × 10^−4^ **Table 2**, **Figure 3A**); this subset consisted 3.1% of total cells from cases compared to 1.7% of cells from controls and achieved an odds ratio of 1.9 (95% CI = 1.3 – 2.7). Conversely, subset 7 (p = 8.8 × 10^−4^, OR = 0.6, 95% CI = 0.5 – 0.8) and subset 12 (p = 2.0 × 10^−3’^, OR = 0.5, 95% CI = 0.4 – 0.8) were underrepresented by RA cells.

**Table 2:** Overview of the subsets found to be significantly expanded in RA. RA proportion reflects the fraction of cells in the subset that were from RA donors. The 95% confidence interval is shown next to odds ratio.

| Condition | Description | Subset | RA Proportion | *P* value | Odds Ratio | Test |
| --- | --- | --- | --- | --- | --- | --- |
| Resting | HLA-DR+,CD27- | 18 | 0.636 | $5.9 \times 10^{-4}$ | 1.9 (1.3 – 2.7) | MASC |
| Stimulated | HLA-DR+, CD27- | 18 | 0.619 | $1.3 \times 10^{-3}$ | 1.7 (1.2 – 2.2) | MASC |
| Resting<br>replication | Gated<br>HLA-DR+, CD27- | NA | NA | $1.1 \times 10^{-2}$ | NA | One-tailed<br>*t* test |

**Figure 3:**
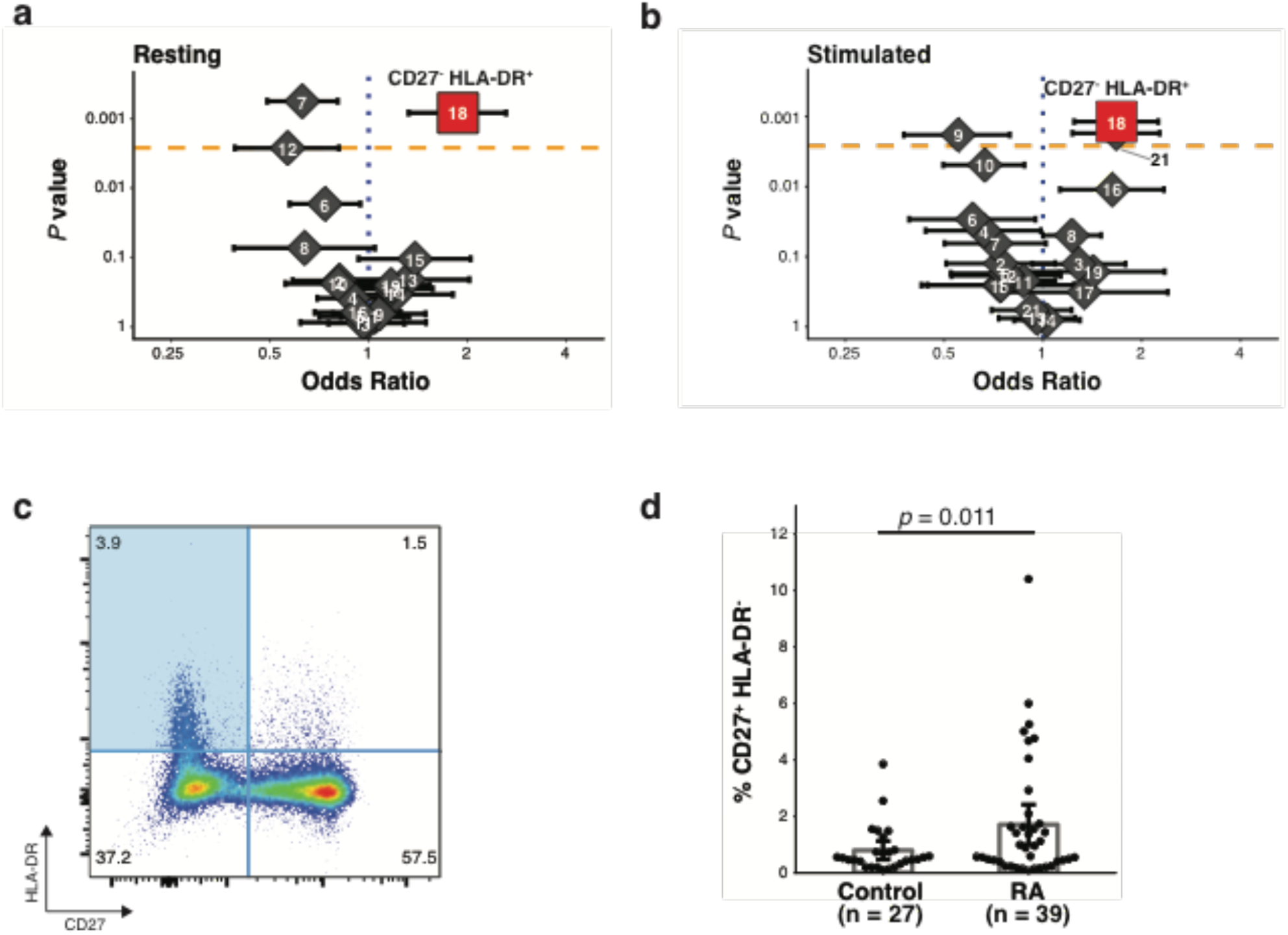
MASC identifies a population that is expanded in RA. (a,b) Odds ratios and association p-values were calculated by MASC for each population identified the resting (a) and stimulated (b) datasets. The yellow line indicates the significance threshold after applying the Bonferroni correction for multiple testing. (c) Flow cytometry dot plot of gated memory CD4+ T cells from a single RA donor shows the gates used to identify CD27^-^ HLA-DR^+^ memory CD4+ T cells (blue quadrant). (d) Flow cytometric quantification of the percentage of CD27^-^, HLA-DR^+^ cells among blood memory CD4+ T cells in an independent cohort of seropositive RA patients (n = 39) and controls (n = 27). Statistical significance was assessed using a one-tailed *t*-test.

To confirm the robustness of this analytical model, we conducted a stringent permutation test that was robust to potential batch effects. For this test, we reassigned case-control labels randomly to each sample; to stringently control for batch effects, retaining the same number of cases and controls in each batch. For each subset we recorded the number of case cells after randomization to define a null distribution. In the real data, we observed that 753 out of the 1184 cells in subset 18 were from case samples. In contrast, when we permuted the data we observed a mean of 550 cells and a standard deviation of 63 cells from case samples. In only 93 out of 100,000 permutations did we observed more than 753 cells from case samples in the randomized data (permutation p = 9.4×10-4, **Supplementary Table 2**). We applied permutation testing to every subset and found that the *p*-values produced by permutation were similar to *p*-values derived from the mixed-effect model framework (Spearman’s r = 0.86, **Figure S1C, D**). When we considered the subsets identified as significant by MASC, both subsets 18 (permutation p = 9.4×10^−4^) and 7 (permutation p = 1.8×10^−4^) retained significance while subset 12 demonstrated nominal evidence of case-control association (permutation p = 1.3×10^−2^).

Next, for each resting subset, we identified corresponding subsets in the stimulated dataset using a cluster centroid alignment strategy to calculate the distance between subsets across datasets (**Methods**). Subset 18 in the resting dataset was most similar to subset 18 in the stimulated dataset, while subset 7 in the resting dataset was most similar to subset 21 in the stimulated dataset (**Figure S1E, F**). Applying MASC, we observed case-control association for subset 18 in the stimulated data (p = 1.2×10^−3^, OR = 1.7, 95% CI = 1.2 – 2.2), while subset 21 (p = 0.55) was not significant in the stimulated data **(Table 2, Figure 3B**). We identified one additional population subset that was significant in the stimulated dataset: subset 20 (p = 1.7×10^−3^, OR = 1.7, 95% CI = 1.2 – 2.3) (**Supplementary Table 3**).

We wanted to ensure that the results we observed were not an artifact of using DensVM to cluster the cytometry data. To this end, we used Phenograph *(23)* and FlowSOM *(31)* to cluster the same dataset. We observed a CD27-, HLA-DR+ cluster with either method (**Figure S3A,C**), which we found to be significantly associated in RA by MASC (**Figure S3B,D**). We also assessed how analysis by MASC compares to Citrus, an algorithm that uses hierarchical clustering to define cellular subsets and then builds a set of models using the clusters to stratify cases and controls. When applied to the same case-control resting dataset, Citrus was unable to produce models with an acceptable cross-validation error rate, regardless of the method used (**Figure S4**).

### CD27- HLA-DR+ CD4+ Effector Memory T Cells Are Expanded in RA Patients

After noting that subset 18 demonstrated robust association to RA, we sought to understand the key features of this population. First, the lack of CD62L indicates that this subset is an effector memory T cell population (**Figure 2E,F**). In order to define the markers that best differentiated this subset from other cells, we calculated the Kullback-Liebler divergence for the expression of each marker between the subset and the full dataset; we applied this approach to both the resting and stimulated datasets (**Methods**, **Figure 4**). The expression of HLA-DR and perforin were notably increased in subset 18 compared to all other cells, while the expression of CD27 was decreased in this subset. Although HLA-DR is a known to be expressed on T cells in response to activation, it takes several days to induce strong HLA-DR expression *(32)*. Thus, it is likely that cells in subset 18 expressed HLA-DR prior to stimulation, such that analyses of both resting and stimulated cells identified the same HLA-DR+ CD27- effector memory T cell population.

**Figure 4:**
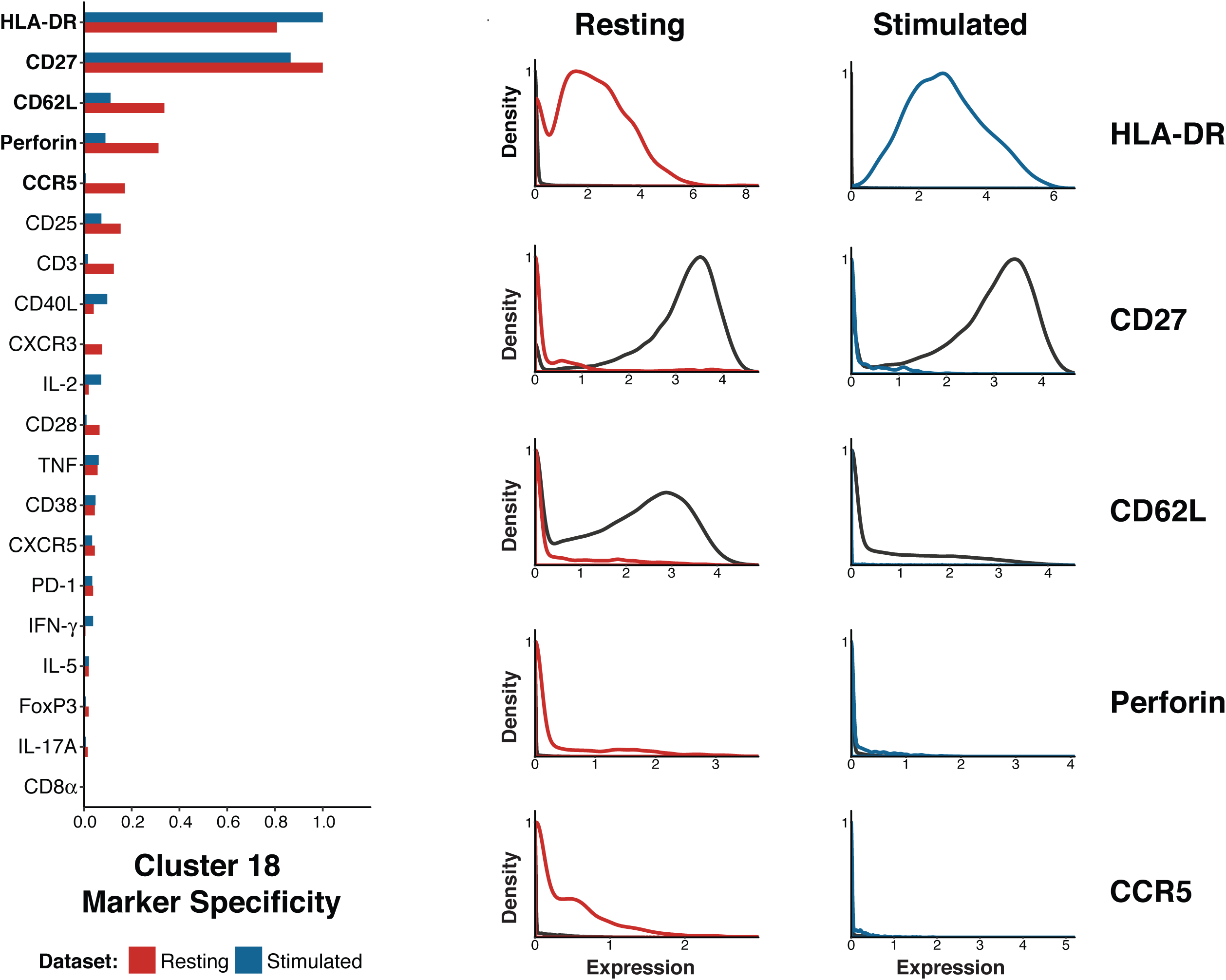
CD27 and HLA-DR expression specifically mark the expanded population. (a) Plot of the Kullback-Liebler divergence for each marker comparing cluster 18 to all other cells in both the resting cell dataset (red) and the stimulated cell dataset (blue). (b) Density plots showing expression of the five markers most different between cluster 18 cells (resting = red, stimulated = blue) and all other cells in the same dataset (black line).

In order to confirm the expansion of the CD27- HLA-DR+ T cell population in RA patients, we evaluated the frequency of CD27- HLA-DR+ T cells in an independent cohort of 39 seropositive RA patients and 27 controls using conventional flow cytometry (**Table 1**). We determined the percentage of memory CD4+ T cells with a CD27- HLA-DR+ phenotype in each group (**Figure 3C**). Consistent with the mass cytometry analysis, CD27- HLA-DR+ cells were significantly expanded in the RA patient samples (p = 0.011, one-tailed *t* test). The frequency of this subset was 0.8% in controls and 1.7% in RA samples, which is similar to the two-fold enrichment we observed in mass cytometry data.

To assess the effect of RA treatment on CD27- HLA-DR+ cell frequency, we quantified CD27- HLA-DR+ cell frequencies in 18 RA patients before and 3 months after initiation of a new medication for RA. In these patients, all of whom experienced a reduction in disease activity after treatment escalation, RA drug therapy significantly reduced the frequency of CD27- HLADR+ cells (**Figure 5A**, p = 0.003, Wilcoxon test). These results confirm that CD27- HLA-DR+ CD4+ T cells are expanded in the circulation of RA patients and decrease with effective disease treatment.

**Figure 5:**
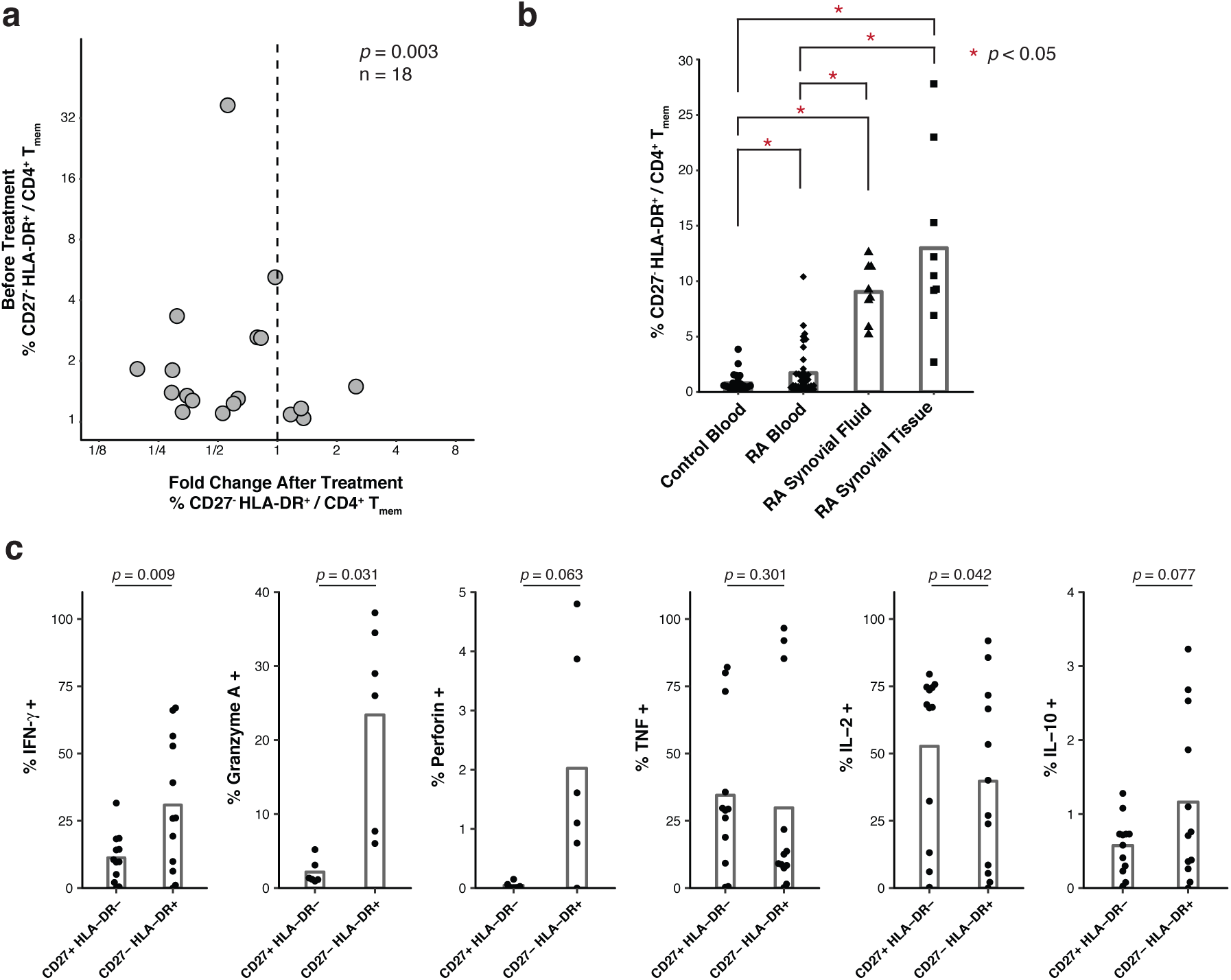
CD27^-^ HLA-DR^+^ memory CD4+ T cells are expanded in the blood and joints of patients with active RA. (a) Flow cytometric quantification of the frequency of CD27- HLA-DR+ memory CD4+ T cells in 18 RA patients prior to starting a new medication, plotted against change in cell frequency after 3 months of new therapy. Treatment significantly reduced CD27- HLA-DR+ cell frequency (p=0.003 by Wilcoxon signed-rank test). b) Flow cytometric quantification of the percentage of memory CD4+ T cells with a CD27- HLA-DR+ phenotype in cells from seropositive RA synovial fluid (n=8) and synovial tissue (n=9), compared to blood samples from RA patients and controls. Blood sample data are the same as shown in panel 3d. Significance was assessed using one-tailed *t*-test and applying a Bonferroni correction for multiple testing. (c) Cytokine expression determined by intracellular cytokine staining of peripheral effector memory CD4+ T cells after *in vitro* stimulation with PMA/ionomycin. The percentage of cells positive for each stain is plotted for CD27+ HLA-DR- and CD27- HLA-DR+ subsets. Each dot represents a separate donor (n = 12; 6 RA patients and 6 controls, except for the quantification of Granyzme A and perforin where n = 6; 3 RA patients and 3 controls).

To determine whether CD27- HLA-DR+ T cells are further enriched at the sites of inflammation in seropositive RA patients, we evaluated T cells in inflamed synovial tissue samples obtained at the time of arthroplasty (**Table 1**) and in inflammatory synovial fluid samples from RA patients. In a set of 9 synovial tissue samples with lymphocytic infiltrates observed by histology, the frequency of CD27- HLA-DR+ cells was increased 5-fold (median 10.5% of memory CD4+ T cells) compared to blood (**Figure 5B**). Notably, in 2 of the tissue samples, >20% of the memory CD4+ T cells displayed this phenotype. CD27- HLA-DR+ cells were similarly expanded in synovial fluid samples from seropositive RA patients (median 8.9% of memory CD4+ T cells, n=8) Thus, CD27- HLA-DR+ T cells are enriched at the primary sites of inflammation in RA patients.

To evaluate the potential function of the CD27- HLA-DR+ cell subset, we assessed production of effector molecules by these cells after *in vitro* stimulation with PMA + ionomycin, a stimulation method that readily reveals T cell capacity for cytokine production. CD27- HLA-DR+ cells produced IFN-γ at a much higher frequency than did CD27+ HLA-DR- cells, which constitute the majority of memory CD4+ T cells (p = 0.009, Wilcoxon signed-rank test, **Figure 5C**). CD27- HLA-DR+ cells also more frequently expressed perforin (p = 0.063, Wilcoxon signed-rank test) and granzyme A (p = 0.031, Wilcoxon signed-rank test), indicating cytolytic capacity. Both T cell populations expressed TNF at the same level (p = 0.301, Wilcoxon signed-rank test), and only slight differences in IL-2 (p = 0.043, Wilcoxon signed-rank test) and IL-10 (p = 0.077, Wilcoxon signed-rank test). Taken together, these observations indicate that the CD27- HLA-DR+ T cells are a Th1-skewed effector memory T cell population capable of producing a range of inflammatory cytokines and cytotoxic molecules.

## Discussion

Mass cytometry has now been successfully applied to decipher heterogeneity of human T cells in multiple settings, including identification of disease-specific changes to circulating immune cell populations and mapping of developmental pathways *(27, 33-36)*. It, and other emerging single cell strategies, offer a promising avenue to characterize the T cell features, and other immunological features, of a wide-range of diseases in humans. However, successful application of mass cytometry on a larger scale (>20 samples) to conduct association studies on clinical phenotypes in humans has been limited thus far *(37-40)*, in part due to limited availability of effective association testing strategies.

Although there are many approaches to cluster single cell data *(19-23, 26, 31)*, extension of these methods to perform case-control association testing is not straightforward. The simplest strategy would be to define subsets of interest by clustering and cells and then comparing frequencies of these subsets in cases and controls with a univariate test such as a t-test or a non-parametric Mann-Whitney test. While commonly used in flow cytometry analysis, this approach is dramatically underpowered, as it relies upon reducing single-cell data to per-sample subset frequencies.

In contrast, our methodology takes full advantage of single cell measurements by using a “reverse-association” framework to test whether case-control status influences the membership of a given cell in a population. That is – each single cell is treated as a single event. But importantly – these events are not independent. Failing to account for dependencies, for example by using a binomial test or not accounting for random effects, results in dramatically inflated statistics and irreproducible associations (**Figure S2**). Using a mixed-effect logistic regression model allows us to account for covariance in single cell data induced by technical and biological factors that could confound association signals (**Figure 1, Figure S2B**), without inflated association tests. MASC also allows users to utilize technical covariates that might be relevant to a single cell measurement (e.g. read depth per cell for single cell RNA-seq or signal quality for mass cytometry) that might influence cluster membership for a single cell.

Here we demonstrate that MASC identified a CD4+ T cell population that is significantly expanded in the circulation of RA patients in two different mass cytometry datasets. We determined a simple marker pair (CD27 and HLA-DR) that allowed for direct, manual gating of the T cell population of interest. We could then interrogate and reproduce the CD27- HLA-DR+ T cell population revealed by MASC using more targeted flow cytometric analysis in blood samples from an independent RA cohort. We note that Citrus, an association strategy to automatically highlight statistical differences between experimental groups and identify predictive populations in mass cytometry data *(41)*, was unable to identify the expanded CD27- HLA-DR+ population. We believe that our methodology compares favorably to Citrus because it incorporates both technical covariates (e.g. batch) and clinical covariates when modeling associations, a key feature when analyzing high-dimensional datasets of large disease cohorts. Additionally, the agglomerative hierarchical clustering framework in Citrus dramatically increases the testing burden and limits power.

The CD27- HLA-DR+ T cells identified by MASC in blood samples from RA patients was further enriched in both synovial tissue and synovial fluid of RA patients. The accumulation of these cells in chronically inflamed RA joints, combined with lack of CD27, suggests that these cells have been chronically activated. Loss of CD27 is characteristic of T cells that have been repeatedly activated, for example T cells that recognize common restimulation antigens *(42-44)*. Importantly, the broader CD27- CD4+ T cell population does not itself differ in frequency between RA patients and controls, in contrast to the expanded population of CD27- HLA-DR+ cells identified by MASC. We hypothesize the CD27- HLA-DR+ T cell population in RA patients may be enriched in RA antigen-specific T cells, offering a potential tool to identify relevant antigens in RA.

Expression of HLA-DR suggests recent or continued activation of these cells *in vivo*. The well-known HLA class II association to RA may also suggest an important role for disease susceptibility for this subset. Despite the suspected chronic activation of these cells, CD27- HLA-DR+ cells do not appear functionally exhausted. CD27- HLA-DR+ cells rapidly produce multiple effector cytokines upon stimulation *in vitro*, with a dramatically increased predisposition to IFN-g production. These cells are also enriched in expression of the cytolytic molecules granzyme A and perforin. CD4+ cytotoxic T cells expressing granzyme A and perforin, also with a CD27- phenotype, are reported to be expanded in patients with chronic viral infections*(45)*. Of note, CD4+ cells that are perforin+ and granzyme A+ have been observed in RA synovial samples*(46, 47)* Interestingly, cytolytic CD4+ T cells lack the capacity to provide B cell help, suggesting that this is a distinct population from the expanded T peripheral helper cell population in seropositive RA*(13, 47)*. Taken together, these findings nominate CD27- HLA-DR+ T cells as a potential pathogenic T cell population that may participate in the chronic autoimmune response in RA *(48)*. Further studies may now evaluate whether this T cell population may have value as a therapeutic target or as a predictive biomarker in RA.

In summary, the MASC single-cell association modeling framework identified a Th1-skewed cytotoxic effector memory CD4+ T cell population expanded in RA using a case-control mass cytometry dataset. The MASC method is extendible to any case-control experiment in which single cell data is available, including flow cytometry, mass cytometry, and single cell RNA-seq datasets. Although current single cell RNA-seq studies are not yet large scale, ongoing projects like the AMP RA/SLE network or the Human Cell Atlas aim to provide these datasets and may benefit from using the MASC framework in case-control testing, or testing for other clinical subphenotypes such as disease progression or activity.

## Materials and Methods

### Experimental Design

Human subjects research was performed in accordance with the Institutional Review Boards at Partners HealthCare and Hospital for Special Surgery via approved protocols with appropriate informed consent as required. Patients with RA fulfilled the ACR 2010 Rheumatoid Arthritis classification criteria, and electronic medical records were used to ascertain patients’ rheumatoid factor and anti-CCP antibody status, C-reactive protein level, and medication use. Synovial tissue samples for mass and flow cytometry were collected from seropositive RA patients undergoing arthroplasty at the Hospital for Special Surgery, New York or at Brigham and Women’s Hospital, Boston. Samples with lymphocytic infiltrates on histology were selected for analysis. Samples inclusion criteria were established prospectively, with the exception of samples that were excluded from the study due to poor acquisition by the mass cytometer.

Synovial fluid samples were obtained as excess material from a separate cohort of patients undergoing diagnostic or therapeutic arthrocentesis of an inflammatory knee effusion as directed by the treating rheumatologist. These samples were de-identified; therefore, additional clinical information was not available.

Blood samples for clinical phenotyping were obtained from consented patients seen at Brigham and Women’s Hospital. We performed medical record review for ACPA (anti-citrullinated protein antibody) positivity according to CCP2 (cyclic citrullinated protein) assays, C-reactive protein (CRP), and RA-specific medications including methotrexate and biologic DMARDS. For blood cell analyses in the cross-sectional cohort, the treating physician measured the clinical disease activity index (CDAI) on the day of sample acquisition. For RA patients followed longitudinally, a new disease-modifying antirheumatic drug (DMARD) was initiated at the discretion of the treating rheumatologist, and CDAIs were determined at each visit by trained research study staff. Blood samples were acquired before initiation of a new biologic DMARD or within 1 week of starting methotrexate and 3 months after initiating DMARD therapy*(49)*. Concurrent prednisone at doses ≤10mg/day were permitted. All synovial fluid and blood samples were subjected to density centrifugation using Ficoll-Hypaque to isolate mononuclear cells, which were cryopreserved for batched analyses.

### Sample Preparation for Mass Cytometry

We rapidly thawed cryopreserved peripheral blood mononuclear cells (PBMCs) at 37° C and isolated total CD4+ memory T cells by negative selection with magnetic bead separation technology. Subsequently, we cultured the CD4+ memory T cells for 24 hours in RPMI 1640, sterile-filtered and supplemented with 15% FBS, 1% Pen/Strep, 0.5% Essential and Non-Essential Amino Acids, 0.5mM Sodium Pyruvate, 30mM HEPES, and 55µM 2-mercaptoethanol. We activated the cells with CD3/CD28 T cell activation beads at a density of 1 bead: 2 cells, and at 6 hours prior to harvesting (t=18 hours), added Monensin and Brefeldin A at a dilution of 1:1000. At 24 hours, we incubated the cells with a rhodium metallointercalator in culture at a final dilution of 1:500 for 15 minutes as a viability measure. We then harvested cells into FACS tubes and washed with CyTOF Staining buffer (CSB) composed of PBS with 0.5% BSA, 0.02% sodium azide, and 2μM EDTA. We spun the cells at 500xg for 7 minutes at room temperature. We incubated the resulting cell pellets with 10ul Fc block and 40ul of CSB for 10 minutes at 4C, and moved to ice until the surface antibody cocktail had been prepared. We then added the CyTOF surface antibodies to the cells and incubated for 30 minutes at 4C on a shaker rack. We washed the cells with 2 ml of CSB and spun at 700xg for 5 minutes at 4C. Post spin, we aspirated the buffer from pellet and added 1 ml of FOXP3 fixative/permeabilization solution supplemented with 1.6% paraformaldehyde. We incubated the cells at room temperature on a gentle shaker in the dark for 45 minutes. We washed the cells with CSB + 0.3% saponin, and spun at 800xg for 5 minutes. To the resulting pellet, we incubated the intracellular antibodies and Iridium in a total volume of 100ul of CSB containing saponin for 35 minutes at r.t. on a gentle shaker. Post-incubation, we washed the cells in PBS and spun at 800xg for 5 minutes. We resuspended the cells in 1 ml of 4% formaldehyde prepared in CyTOF PBS and incubated for 10 minutes at room temperature on a gentle shaker, followed by another PBS wash and spin at 800xg for 5 minutes. We washed the pellet in deionized water, spun at 700xg for 6 minutes, and subsequently resuspended the resulting pellet in deionized water at 700,000 cells per ml for analysis via the CyTOF 2. We transferred the suspensions to new FACS tubes through a 70μm cell strainer and added 4-element MaxPar beads at a ratio of 1:10 by volume prior to acquisition.

### Mass Cytometry Panel Design

We designed an antibody panel for mass cytometry with the goal of both accurately identifying CD4+ effector memory T cell populations and measuring cellular heterogeneity within these populations. We chose markers that fell into one of five categories to generate a broadly informative panel: chemokine receptors, transcription factors, lineage markers, effector molecules, and markers of cellular activation and exhaustion (**Supplemental Table 1**). Antibodies were obtained from DVS or from the Longwood Medical Area CyTOF Antibody Resource Core (Boston, MA).

### Mass Cytometry Data Acquisition

We analyzed samples at a concentration of 700,000 cells/ml on a Fluidigm-DVS CyTOF 2 mass cytometer. We added Max Par 4-Element EQ calibration beads to every sample that was run on the CyTOF 2, which allowed us to normalize variability in detector sensitivity for samples run in different batches using previously described methods*(50)*. We used staining for iridium and rhodium metallointercalators to identify viable singlet events. We excluded samples where acquisition failed or only yielded a fraction of the input cells (< 5%). The criteria for sample exclusion were not set prospectively but were maintained during all data collection runs.

Since samples were processed and analyzed on different dates, we ran equal numbers of cases and controls each time to guard against batch effects. However, as CyTOF data are very sensitive to day-to-day variability, we took extra steps to pre-process and normalize data across the entire study.

### Flow Cytometry Sample Preparation

For flow cytometry analysis of the validation and longitudinal blood cohorts and synovial samples, cryopreserved cells were thawed into warm RPMI/10% FBS, washed once in cold PBS, and stained in PBS/1% BSA with the following antibodies for 45 minutes: anti-CD27-FITC (TB01), anti-CXCR3-PE (CEW33D), anti-CD4-PE-Cy7 (RPA-T4), anti-ICOS-PerCP-Cy5.5 (ISA-3), anti-CXCR5-BV421 (J252D4), anti-CD45RA-BV510 (HI100), anti-HLA-DR-BV605 (G46-6), anti-CD49d-BV711 (9F10), anti-PD-1-APC (EH12.2H7), anti-CD3-AlexaFluor700 (HIT3A), anti-CD29-APC-Cy7 (TS2/16), propidium iodide. Cells were washed in cold PBS, passed through a 70-micron filter, and data acquired on a BD FACSAria Fusion or BD Fortessa using FACSDiva software. Samples were analyzed in uniformly processed batches containing both cases and controls.

### Flow Cytometry Intracellular Cytokine Staining

Effector memory CD4^+^ T cells were purified from cryopreserved PBMCs by magnetic negative selection (Miltenyi) and rested overnight in RPMI/10%FBS media. The following day, cells were stimulated with PMA (50ng/mL) and ionomycin (1μg/mL) for 6 hours. Brefeldin A and monensin (both 1:1000, eBioscience) were added for the last 5 hours. Cells were washed twice in cold PBS, incubated for 30 minutes with Fixable Viability Dye eFluor 780 (eBioscience), washed in PBS/1%BSA, and stained with anti-CD4-BV650 (RPA-T4), anti-CD27-BV510 (TB01), anti-HLA-DR-BV605 (G46-6), anti-CD20-APC-Cy7 (2H7), and anti-CD14-APC-Cy7 (M5E2). Cells were then washed and fixed and permeabilized using the eBioscience Transcription Factor Fix/Perm Buffer. Cells were then washed in PBS/1%BSA/0.3% saponin and incubated with anti-IFN-γ-FITC (B27), anti-TNF-PerCp/Cy5.5 (mAb11), anti-IL-10-PE (JES3-9D7) and anti-IL-2-PE/Cy7 (MQ1-17H12) or anti-granzyme A-AF647 (CB9) and anti-perforin-PE/Cy7 (B-D48) for 30 minutes, washed once, filtered, and data acquired on a BD Fortessa analyzer. Gates were drawn to identify singlet T cells by FSC/SSC characteristics, and dead cells and any contaminating monocytes and B cells were excluded by gating out eFluor 780-positive, CD20+, and CD14+ events.

### Synovial Tissue Processing

Synovial samples were acquired after removal as part of standard of care during arthroplasty surgery. Synovial tissue was isolated by careful dissection, minced, and digested with 100μg/mL LiberaseTL and 100μg/mL DNaseI (both Roche) in RPMI (Life Technologies) for 15 minutes, inverting every 5 minutes. Cells were passed through a 70μm cell strainer, washed, subjected to red blood cell lysis, and cryopreserved in Cryostor CS10 (BioLife Solutions) for batched analyses.

## Statistical Analysis

### Mixed effects modeling of Associations of Single Cells (MASC)

Here, we present a flexible method of finding significant associations between subset abundance and case-control status that we have named MASC (Mixed effects modeling of Associations of Single Cells). The MASC framework has three steps: (1) stringent quality control, (2) definition of population clusters, and (3) association testing. Here we assume that we have single cell assays each quantifying *M* possible markers where markers can be genes (RNA-seq) or proteins (cytometry).

#### Quality control

To mitigate the influence of batch effects and spurious clusters, we first remove poorly recorded events and low quality markers before further analysis. We remove those markers (1) that have little expression, as these markers are not informative, and (2) with significant batch variability. First, we concatenated samples by batch and measured the fraction of cells negative and positive for each marker. We then calculated the ratio of between-batch variance to total variance for each marker’s negative and positive populations, allowing us to rank and retain 20 markers that were the least variable between batches. We also removed markers that were either uniformly negative or positive across batches, as this indicated that the antibody for that marker was not binding specifically to its target. For single-cell transcriptomic data, an analogous step would involve removing genes with low numbers of supporting reads or genes whose expression varies widely between batches.

Once low quality markers have been identified and removed, we remove events that are likely to be artifacts. We first remove events that have extremely high signal for a single marker: events that have recorded expression values at or above the 99.9^th^ percentile for that marker are removed. These events are unlikely to be intact, viable cells. Next, a composite “information content” score (eq 1) for each event *i* is created in the following manner: the expression *x* for each marker *M* is rescaled from 0 to 1 across the entire dataset to create normalized expression values *y*_*i*_ for each event *i*. The sum of these normalized expression values is used to create the event’s information content score.

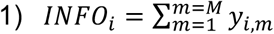

The information content score reflects that events with little to no expression in every channel are less informative than events that have more recorded expression. Events with low scores (*INFO*_*i*_ < 0.05) and are unlikely to be informative in downstream analysis are removed. In addition, events that derive more than half of their information content score from expression in a single channel are also removed (2):

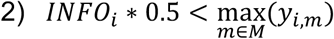

Potential explanations for these events include poorly stained cells or artifacts caused by the clumping of antibodies with DNA fragments. A final filtering step retains events that were recorded as having detectable expression in at least *M*_*min*_ markers, where *M*_*min*_ may vary from experiment to experiment based on the panel design and expected level of co-expression between channels. The quality control steps described here are specific for mass cytometry analysis and will need to be optimized for use with transcriptomic data.

#### Clustering

After applying quality control measures to each sample, we combine data from cases and controls into a single dataset. It is critical to ensure that each sample contributes an equal numbers of cells to this dataset, as otherwise the largest samples will dominate the analysis and confound association testing. We suggest sampling an equal number of cells from each sample. After sampling the data, we partition these cells into populations using DensVM, which performs unsupervised clustering based on marker expression. We note that partitioning the data can be accomplished with different clustering approaches – such as SPADE or PhenoGraph for mass cytometry data – or even by using traditional bivariate gating, as MASC is not dependent on any particular method of clustering (**Figure S3**)

#### Association testing

Once all cells have been assigned to a given cluster, the relationship between single cells and clusters is modeled using mixed-effect logistic regression to account for donor or technical variation (eq 3). We model the age and sex of sample *k as* fixed effect covariates, while the donor and batch that cell *i* belongs to are modeled as random effects. The random effects variance-covariance matrix treats each sample and batch as independent gaussians. Each cluster is individually modeled. Note that this baseline model does not explicitly include any single cell expression measures.

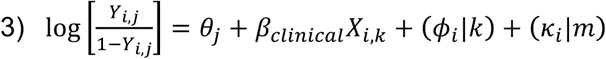

Where *Y*_*i*_ is the odds of cell *i* belonging to cluster *j, θ*_*i*_ is the intercept for cluster *j, β*_*clinical*_ is a vector of clinical covariates for the k^th^ sample, (*ϕ*_*i*_|*k*) is the random effect for cell *i* from k^th^ sample, (*κ*_*i*_|*m*) is the random effect for cell *i* from batch *m*.

To determine if any clusters are associated with case-control status, we include an additional covariate that indicates whether the k^th^ sample is a case or control (eq. 4)

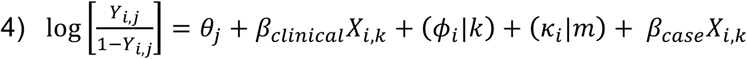

Here, *Y*_*i,j*_ is the odds of cell *i* belonging to cluster *j*, *θ*_*i*_ is the intercept for cluster *j, β*_*clinical*_ is a vector of clinical covariates for the k^th^ sample, *ϕ_i_*|*k* is the random effect for cell *i* from k^th^ sample, (*k*_*i*_|*m*) is the random effect for cell *i* from batch *m, β*_*case*_ indicates the effect of k^th^ sample’s case-control status.

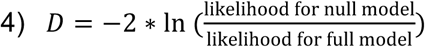

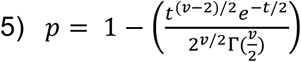

We compare the two models using a likelihood ratio test (eq. 5) to find the test statistic *D*, which is the ratio of the likelihoods for the baseline and full models. The term *D* is distributed under the null by a *χ*^2^ distribution with 1 degree of freedom, as there is only one additional parameter in the full model compared to the null (case-control status). We derive a *p*-value by comparing test statistic *D* of the likelihood ratio test to the value of the *χ*^2^ distribution with 1 degrees of freedom (eq. 6), allowing us to find clusters in which case-control status significantly improves model fit. A significant result indicates that cluster membership for a single cell is influenced by case-status after accounting for technical and clinical covariates. The effect size of the case-control association can be estimated by calculating the odds ratio from *β*_*case*_. If a dataset includes multiple groups, then we can test for association between *g* groups using *g*-1 indicator variables. This approach allows us to capture inter-individual differences between donors, as well as model the influence of technical and clinical covariates that might influence a cell to be included as a member of one cluster versus another.

### Mass Cytometry Data Analysis

We analyzed 50 samples (26 cases, 24 controls) in the resting condition and 52 samples (26 cases, 26 controls) in the stimulated condition by CyTOF. Two samples were only analyzed after stimulation due to low numbers of available PBMCs. We ran aliquots of a standard PBMC sample alongside cases and controls with each CyTOF run to allow us to measure batch variability directly, as these aliquots should not be biologically dissimilar. We then used these data to find markers that had stained poorly or varied significantly between batches and removed them analysis. After acquisition, each sample was gated to a CD4^+^, CD45RO^+^ population using FlowJo 10.1 (TreeStar, USA) and combined into a single dataset before analyzing the data using MASC as previously described. We performed the data filtration steps requiring cells to demonstrate expression in at least 5 markers (*M*_*min*_ = 5). This removed 3.0- 6.0% of all events captured in each sample. We first removed the initial noise factor applied to all zero expression values in mass cytometry by subtracting 1 from expression values and setting any negative values to 0, then applied the inverse hyperbolic sine transform with a cofactor of 5 to the raw expression data, using the following equation: 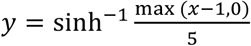

To partition the data, we first randomly selected 1000 cells from each sample and applied the t-Distributed Stochastic Neighbor Embedding (tSNE) algorithm (Barnes-Hut implementation*(51)*) to the reduced dataset with the following parameters: perplexity = 30 and theta = 0.5. We did not include channels for CD4 or CD45RO in the SNE clustering as these markers were used for gating samples. To identify high-dimensional populations, we used a modified version of DensVM*(26)*. DensVM performs kernel density estimation across the dimensionally reduced SNE map to build a training set, then assigns cells to clusters by their expression of all markers using an SVM classifier. We modified the DensVM code to increase the range of potential bandwidths searched during the density estimation step and to return the SVM model generated from the SNE projection.

Association testing for each cluster was performed using mixed effect logistic regression according to the MASC method. Donor and batch were included as random-effect covariates, donor sex and age was included as a fixed-effect covariate, and donors were labeled as either cases (RA) or controls (OA). To confirm the associations found by MASC, we conducted exact permutation testing in which we permuted the association between case-control status and samples within batches 10,000 times, measuring the fraction of cells from RA samples that contributed to each cluster in each permutation. This allowed us to build an empirical null for the case-control skew of each cluster, and we could then determine for each cluster how often a skew equal or greater to the observed skew occurred. We adjusted the p-values for the number of tests we performed (the number of clusters analyzed in each condition) using the Bonferroni correction.

We aligned subsets between experiments using the following strategy: In each experiment, expression data was first scaled to mean zero, variance one to account for differences in sensitivity. We used the mean expression value for marker column to define a centroid for each cluster. We then aligned clusters by calculating the Euclidean distance between cluster centroids in the query and target datasets.

We wanted to determine which markers best separated a given population from the rest of the data in a quantitative manner, as finding a set of population-specific markers is crucial for isolating the population *in vivo*. In order to do this, we examined the distribution of expression for each marker individually in the entire dataset (*Q*) and the population of interest (*P*). We then grouped expression values for *P* and *Q* into 100 bins, normalized the binned vector to 1, and calculated the Kullback-Leibler divergence from *Q* to *P* for that marker with the following equation:

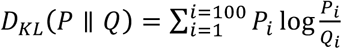

The divergence score can be interpreted as a measure of how much the distribution of expression for a given marker in the entire dataset resembles the distribution of expression for that marker in the population of interest. Higher scores represent lower similarity of the marker’s expression distributions for *P and Q*, indicating that the expression profile of that marker is more specific for that population. By calculating this score for every marker, we can rank and identify markers that best differentiate the population of interest from the dataset.

We independently clustered the resting dataset with Phenograph and FlowSOM using the same cells and markers used to cluster the data with DensVM. We set *k* to 19 for FlowSOM clustering to match the number of clusters found by DensVM; for Phenograph, we used the default setting of k = 30.

All analyses were performed using custom scripts for R 3.4.0^48^. We used the following packages: *flowCore*^49^ to read and process FCS files for further analysis, *lme4*^50^ to apply mixed effect logistic regression, *ggplot2*^51^ and *pheatmap*^52^ for data visualization, and *cytofkit*^53^ to implement FlowSOM and Phenograph.

### Flow Cytometry Data Analysis

Flow cytometry data were analyzed using FlowJo 10.0.7, with serial gates drawn to identify singlet lymphocytes by FSC/SSC characteristics. Viable memory CD4+ T cells were identified as propidium iodide-negative CD3+ CD4+ CD45RA-cells. We then calculated the frequency of HLA-DR-CD27+ cells among memory CD4+ T cells.

## List of Supplementary Materials

Figure S1. Association Permutation Testing and Cluster Alignment.

Figure S2. MASC Type 1 Error.

Figure S3. Clustering with Phenograph and FlowSOM.

Figure S4. Association Testing with Citrus.

Table S1. Panel design for mass cytometry experiments.

Table S2. MASC analysis of the 19 clusters identified in the resting dataset.

Table S3. MASC analysis of the 21 clusters identified in the stimulated dataset.

## Acknowledgements

This work was supported in part by funding from the National Institutes of Health (UH2AR067677, U19AI111224, and 1R01AR063759 (to S.R.)), the Doris Duke Charitable Foundation Grant #2013097, and funding from the Smith Fund at Brigham and Women’s Hospital, T32 AR007530-31 and the William Docken Inflammatory Autoimmune Disease Fund (to M.B.B), Rheumatology Research Foundation Tobe and Stephen Malawista, MD Endowment in Academic Rheumatology (to D.A.R), R01 AR064850-03 (to Y.C.L).

## Author Contributions

C.Y.F., S.R. conceived the MASC association testing method. C.Y.F., S.R., and K.F.S. conducted all statistical analyses. D.A.R., N.C.T., S.K.H., M.F.G., J.E., M.B.B., and S.R. designed and conducted all mass cytometry, flow cytometry, and functional immunology assays. D.A.R., L.T.D., M.E.W., E.M.M., J.S.C., S.M.H., D.J.T., V.P.B., E.W.K., and Y.C.L. recruited patients and obtained samples for this study. C.Y.F, D.A.R. and S.R. wrote the initial manuscript. All authors edited and revised the final manuscript.

## Data and Software Availability

The data that support the findings of this study are available from the corresponding author upon reasonable request. MASC and all other custom scripts used in this analysis are available from the corresponding author upon request.

